# SpNeigh: spatial neighborhood and differential expression analysis for high-resolution spatial transcriptomics

**DOI:** 10.1101/2025.11.07.687304

**Authors:** Jinming Cheng, Pierce K.H. Chow, Nan Liu

## Abstract

Spatial transcriptomics technologies such as Xenium, MERFISH, and Visium HD enable high-resolution profiling of gene expression while preserving tissue architecture. However, most computational methods for spatial analysis do not explicitly model local tissue context, such as boundaries, neighborhoods, or gradients. Here we present SpNeigh, an R package for spatial neighborhood analysis and spatially-aware differential expression modeling. SpNeigh includes tools for boundary detection, spatial neighborhood extraction, distance-based weighting, and gradient-based statistical testing. It supports both region-based differential expression and smooth spatial modeling using spline-based regression, along with a spatial enrichment index that identifies genes enriched near defined spatial features. We demonstrate the utility of SpNeigh across multiple platforms and tissues, including mouse brain, human breast cancer, and human liver, revealing intermediate populations at tissue interfaces, immune microenvironment differences, and spatially zonated gene expression patterns. SpNeigh offers a flexible and interpretable framework for dissecting spatial gene expression dynamics in complex tissues.

## 1 Introduction

Spatial transcriptomics technologies such as 10x Genomics Xenium [1], Vizgen MERFISH (MERSCOPE) [2], and 10x Visium HD [3] enable high-throughput measurement of spatially resolved gene expression across hundreds to thousands of genes, preserving spatial context within tissues. These platforms provide rich opportunities to investigate cellular organization, tissue structure, and microenvironmental interactions at unprecedented resolution. To analyze such data, a growing number of computational frameworks have been developed, including general-purpose toolkits such as *Giotto* [4] and *Squidpy* [5], which support spatial clustering, neighborhood graph construction, visualization, and exploratory analysis.

While these frameworks offer broad analytical capabilities, many spatial transcriptomics analyses ultimately rely on differential expression (DE) to detect transcriptional differences across spatial contexts. Existing DE methods for spatial data, such as *SpatialDE* [6] and *SPARK* [7], are primarily designed for the global detection of spatially variable genes (SVGs) using Gaussian process regression or spatial auto-correlation. However, they do not explicitly model local spatial contexts—such as boundaries between regions or proximity to specific tissue structures—which are critical for understanding tissue interfaces, transitional zones, and niche-specific gene regulation.

To address this gap, we developed *SpNeigh*, an R package that provides a spatially informed neighborhood analysis framework tailored for high-resolution spatial transcriptomics data. *SpNeigh* introduces a novel design that leverages boundary-aware and centroid-aware spatial weights to quantify local context, enabling downstream analyses such as differential expression between boundary-adjacent and interior cells, gradient-based modeling of gene expression along spatial gradients, and spatial enrichment scoring. The package integrates multiple components—including spatial boundary detection, neighborhood interaction analysis, and spatial weight computation—into a unified and reproducible workflow, with built-in visualization and statistical testing functionality.

*SpNeigh* supports data from multiple spatial transcriptomics platforms, including Xenium, MERFISH, and Visium HD, and can be flexibly applied across a range of tissues and biological contexts. By modeling local spatial neighborhoods and continuous gradients, *SpNeigh* complements existing tools and provides a unified framework for dissecting region-specific transcriptional variation.

We demonstrate the utility of *SpNeigh* across diverse datasets, including mouse brain, human breast cancer, and healthy human liver. In each case, *SpNeigh* reveals spatially enriched genes and local transcriptional gradients that are not easily captured by global or unsupervised methods. Together, these results highlight the value of neighborhood-aware spatial modeling and establish *SpNeigh* as a versatile toolkit for spatial expression analysis.

## 2 Results

### 2.1 Overview of the SpNeigh framework

To characterize local tissue architecture and spatial gene expression patterns, we developed *SpNeigh*, a modular R package for spatial neighborhood analysis and spatially-aware differential expression analysis. The *SpNeigh* workflow consists of five core components (Fig. 1): (i) data input, (ii) boundary detection and neighborhood construction, (iii) spatial weighting, (iv) neighborhood interaction analysis, and (v) spatial differential and enrichment analysis.

**Fig. 1.**
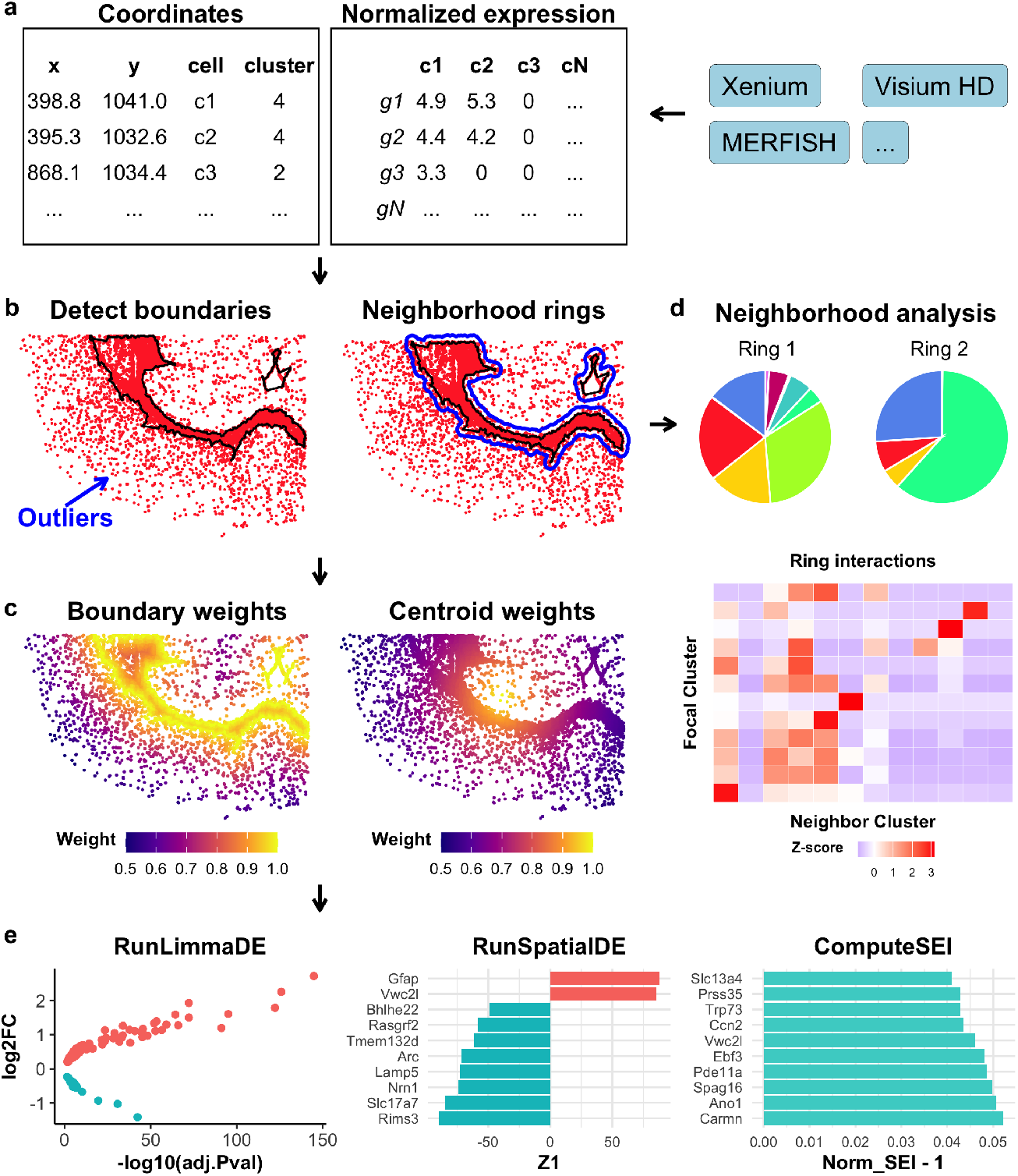
Overview of the SpNeigh workflow. **a**. Input includes a spatial coordinate data frame (x, y, cell, cluster) and a normalized expression matrix. Data can originate from platforms such as Xenium, Visium HD, MERFISH, or others. **b**. Spatial boundary detection and neighborhood extraction. Left: Cluster boundaries are identified after removing spatial outliers based on local k-nearest neighbor density. Right: Ring regions are constructed by buffering outward from the cluster boundaries. Black lines denote cluster boundaries; blue lines indicate outer ring boundaries. **c**. Spatial weight computation. Cells are assigned weights based on their distance to either the boundary (left) or the centroid (right) of the cluster using inverse distance decay. Weights range from 0 (far) to 1 (close), reflecting proximity. **d**. Neighborhood composition and interaction analysis. Top: Pie chart showing the proportion of neighboring cell types within the rings. Bottom: Heatmap of spatial interaction scores between focal and neighboring clusters. **e**. Downstream analyses enabled by SpNeigh. Left: Differential expression analysis between cells of the same cluster in the inner region versus the ring. Middle: Spatial differential expression analysis using smooth functions of distance-based weights. Right: Spatial enrichment analysis quantifying expression bias relative to spatial proximity.

#### (i) Input data

*SpNeigh* operates on a coordinate data frame containing the `x, y, cell,` and `cluster` columns, together with a normalized gene expression matrix or `dgCMatrix`. These inputs can be extracted from commonly used spatial transcriptomics platforms, including 10x Genomics Xenium, Vizgen MERFISH, and 10x Visium HD.

#### (ii) Boundary detection and neighborhood construction

Spatial boundaries are identified for a specified cell population using clustering and density-based outlier removal. Optional buffer regions can be constructed around these boundaries to define concentric neighborhood rings, enabling comparative analysis between intra-region and peri-region cells.

#### (iii) Spatial weighting

For cells within a region, *SpNeigh* computes spatial weights based on distance to either the region boundary or the population centroid. These weights are scaled between 0 and 1, with higher values indicating closer proximity to the spatial feature of interest. Multiple decay schemes (e.g., inverse, Gaussian) are supported.

#### (iv) Neighborhood analysis

Cells located in neighborhood rings are extracted, and their composition is visualized using pie charts and bar plots. A spatial interaction matrix is computed by counting the *k* nearest neighbors for each focal cluster and tabulating the frequencies of neighboring cluster identities.

#### (v) Downstream spatial modeling

*SpNeigh* supports differential expression testing between cells inside and outside a region via `RunLimmaDE`, which leverages the widely used *limma* framework for linear modeling and empirical Bayes moderation [8]. Spatial expression gradients are modeled using `RunSpatialDE`, which fits gene expression as a smooth function of spatial weights using a spline-based linear model, also implemented within the *limma* framework. In parallel, spatial enrichment is quantified using the Spatial Enrichment Index (SEI), which measures the weighted average expression of each gene relative to spatial proximity.

Together, these components provide a flexible and interpretable framework for dissecting fine-grained spatial organization and gene expression dynamics in diverse tissues and technologies. For convenience, several functions such as boundary detection and spatial plotting support direct input from `Seurat` objects, though core modeling functions require explicit expression matrices and spatial weights.

### 2.2 SpNeigh reveals intermediate cell populations at brain region boundaries in mouse cortex

The mouse brain is anatomically well-characterized and widely used for benchmarking spatial transcriptomics tools, with extensive references available for cell type classification and regional annotations [9–11]. To demonstrate the utility of boundary-aware analysis in *SpNeigh*, we analyzed a publicly available mouse brain Xenium dataset (tiny version) from 10x Genomics. After quality control filtering to remove low-quality cells (total count or gene count thresholds), 36,402 high-quality cells remained out of 36,602 total. The *Seurat* [12] pipeline was applied for preprocessing and clustering, resulting in 12 clusters at a resolution of 0.1. These clusters were manually annotated based on known brain anatomical regions. To refine cellular identity, we also performed cell-level annotation using *SingleR* [13], referencing a curated subset of the large-scale mouse brain single-cell RNA sequencing (scRNA-seq) dataset from Yao et al. [14]. The resulting *SingleR* annotations (Supplementary Fig. 1a) were manually merged based on known taxonomy to reduce redundancy (see Supplementary Table 1). The clustering, region-level, and cell-level annotations are shown in Fig. 2a.

**Fig. 2.**
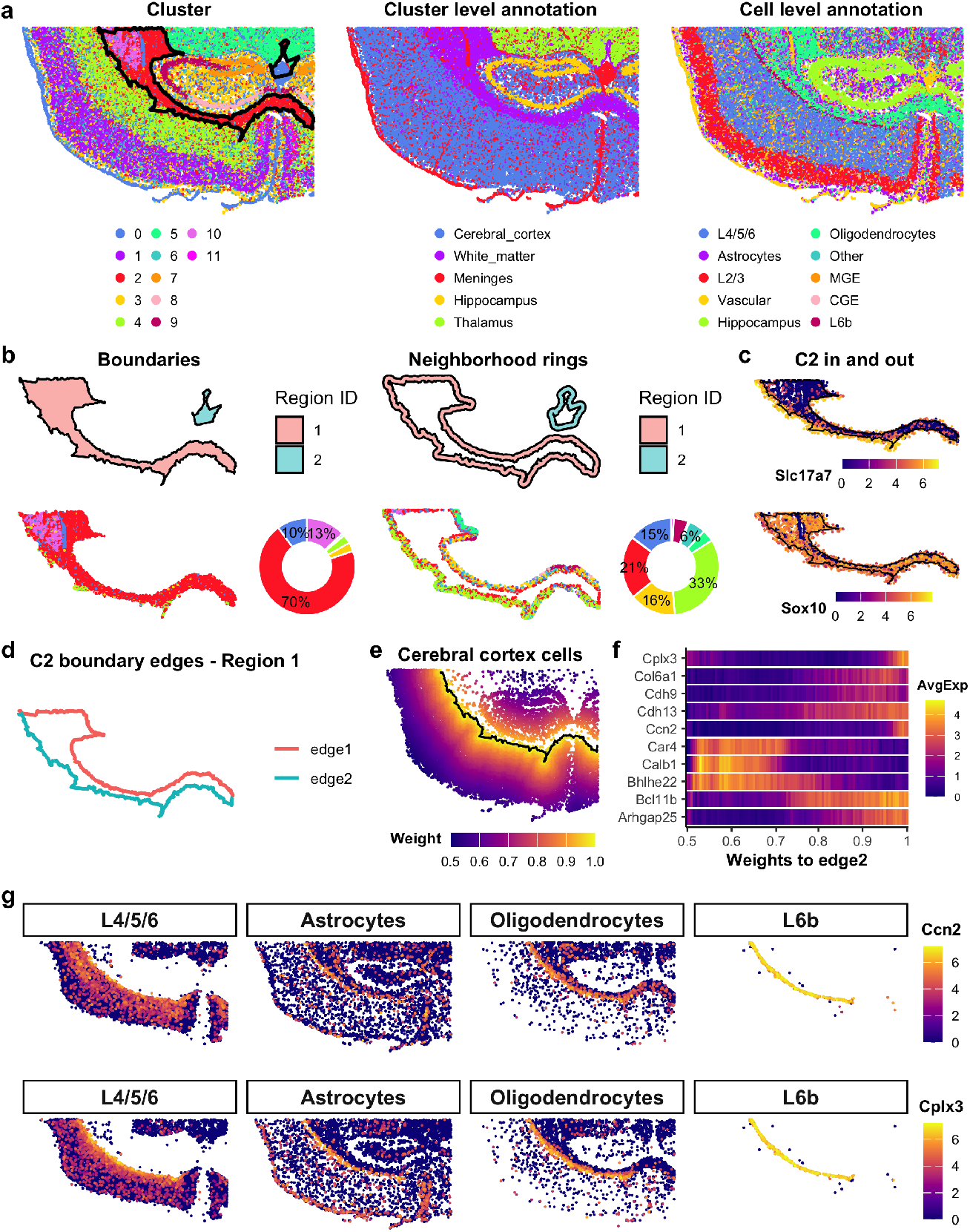
SpNeigh reveals intermediate cell populations near boundaries in mouse brain Xenium data. **a**. Spatial plots showing different annotation types. Left: Cells colored by clusters with overlaid boundaries of cluster 2. Middle: Manual cluster-level annotations based on brain anatomy. Right: Reference-based single-cell annotations, with selected subtypes merged. CGE: caudal ganglionic eminence; MGE: medial ganglionic eminence. **b**. Neighborhood analysis of cluster 2. Top: Boundary and ring regions. Bottom: Cells within boundary and ring regions for region 1, with donut plots showing cluster proportions (labels shown for proportions ≥ 5%). **c**. Expression of Slc17a7 and Sox10 in cluster 2 cells inside boundaries and surrounding rings. Slc17a7, a marker of cortical excitatory neurons, shows elevated expression in outer cells near the boundary. Sox10 is broadly expressed in oligodendrocytes and remains consistent across both inner and outer cells in cluster 2. **d**. Boundary 1 of cluster 2 split into discrete edges. **e**. Spatial weights relative to edge 2 for cortical cells. Black line indicates edge 2. **f**. Top spatially varying genes identified by RunSpatialDE using weights from edge 2. **g**. Expression of Ccn2 and Cplx3 near edge 2. Cells include cortical layer 4/5/6 neurons, L6b neurons, astrocytes, and oligodendrocytes. L6b cells are localized along edge 2.

We focused our initial analysis on cluster 2, corresponding to the white matter region dominated by oligodendrocytes. This area connects the outer cerebral cortex to deeper brain regions and lies between the cortex and hippocampus. Using *SpNeigh*, two spatial boundaries and their corresponding neighborhood ring regions were detected for cluster 2. Cells within the primary (largest) boundary and surrounding ring were extracted for comparison (Fig. 2b). Inside the boundary, 70% of cells belonged to cluster 2, with clusters 10 and 0 contributing 13% and 10%, respectively. In contrast, within the neighborhood ring, cluster 4 accounted for 33% of cells, followed by clusters 2 (21%), 3 (16%), and 0 (15%). These distinct neighborhood compositions suggest that cluster 2 cells located within the boundary and in the surrounding ring may reside in different microenvironments.

To investigate this difference, we applied `RunLimmaDE` to compare cluster 2 cells inside the boundary versus those in the ring. We found that the cortical marker gene *Slc17a7* was highly expressed in cluster 2 cells within the neighborhood ring but nearly absent within the boundary (Fig. 2c; Supplementary Fig. 1b). In contrast, the oligodendrocyte marker gene *Sox10* was consistently expressed in both groups. This suggests that cluster 2 cells in the ring may represent intermediate cells at the interface between cortical neurons and white matter oligodendrocytes.

Given the layered architecture of the cortex, we next explored expression gradients in cerebral cortex cells adjacent to the white matter. The primary boundary of cluster was split into two directional edges; edge 2 delineates the boundary between cortex and white matter (Fig. 2d). Spatial weights relative to edge 2 were computed and used as input for gradient modeling with `RunSpatialDE` (Fig. 2e). A heatmap of the top 10 spatially varying genes showed distinct expression trends along this axis (Fig. 2f). Notably, *Ccn2* (also known as *Ctgf*) and *Cplx3* were enriched near edge 2, consistent with prior studies identifying these genes as markers of layer 6b (L6b)—the deepest cortical layer [14–17]. Both genes exhibited strong expression in L6b cells, with intermediate levels in neighboring L4/5/6 neurons, astrocytes, and oligodendrocytes (Fig. 2g; Supplementary Fig. 1c), further supporting the presence of transitional or mixed-identity cells near the white matter boundary.

These findings highlight the ability of *SpNeigh* to detect spatial gene expression gradients and reveal boundary-associated intermediate populations that may otherwise be overlooked. This boundary-aware modeling is especially valuable for understanding the fine-grained organization and developmental transitions in complex tissues such as the brain.

### 2.3 Benchmarking differential expression and spatial modeling methods in SpNeigh

To evaluate the performance of the DE function `RunLimmaDE` in *SpNeigh*, we benchmarked it against the widely used `FindAllMarkers` function in *Seurat* using the mouse brain Xenium dataset. Both methods were applied to identify marker genes for five major cell types based on cluster-level annotations. `RunLimmaDE` completed in approximately 2 seconds, more than 10 times faster than `FindAllMarkers`, which required 26 seconds (Fig. 3a). The top marker genes identified by `RunLimmaDE` clearly separated distinct cell types, as shown in the dot plot of average expression and percent detection (Fig. 3b). DE results from both methods were highly concordant, with an overall Spearman correlation of 0.81 between log_2_ fold changes across all cell types (Fig. 3c).

**Fig. 3.**
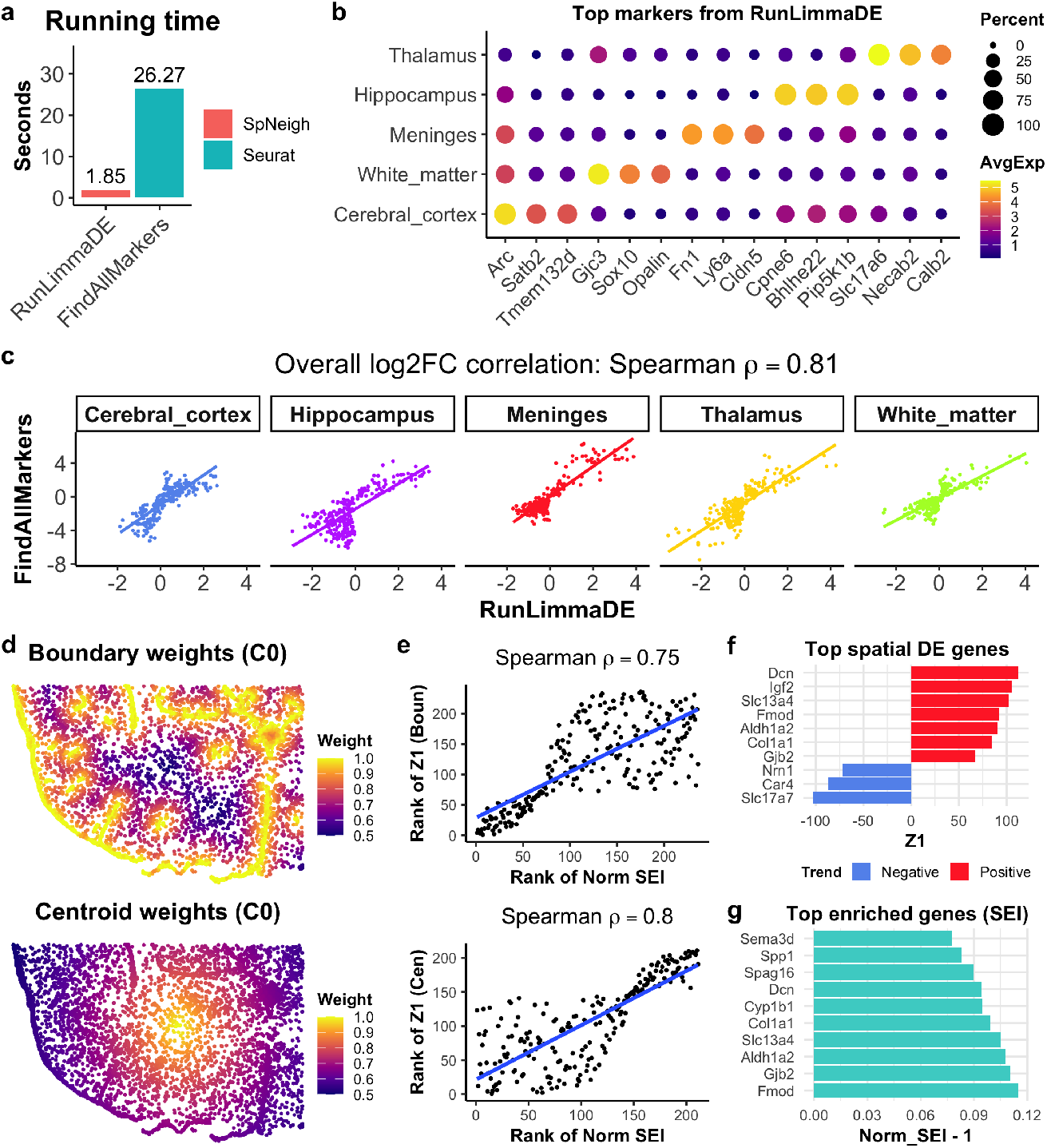
Benchmarking and comparative evaluation of differential expression and spatial enrichment methods in SpNeigh. **a**. Comparison of runtimes for marker gene identification using RunLimmaDE (SpNeigh) and FindAllMarkers (Seurat) across all cell types defined by cluster-level annotation. **b**. Dot plot of top marker genes identified by RunLimmaDE for each annotated cell type. **c**. Scatter plots comparing log2 fold changes (log2FC) between RunLimmaDE and FindAllMarkers for each cell type. The overall Spearman correlation is 0.81. **d**. Spatial weight distributions for cells in cluster 0 based on proximity to the cluster’s boundary (left) and centroid (right). **e**. Scatter plots comparing the gene-wise ranks of Z1 coefficients from RunSpatialDE and normalized spatial enrichment index (SEI) values derived from boundary and centroid weights. Spearman correlations are 0.75 (boundary) and 0.80 (centroid). **f**. Top spatially varying genes identified by RunSpatialDE using boundary-derived weights. A positive Z1 indicates increasing expression toward the boundary; a negative Z1 indicates decreasing expression. **g**. Top spatially enriched genes ranked by normalized SEI from boundary weights. The x-axis shows normalized SEI minus 1 for visualization.

The spatial modeling methods implemented in *SpNeigh*—`RunSpatialDE` and `ComputeSpatialEnrichmentIndex`—are not directly comparable to tools such as *SpatialDE* or *SPARK*, which are designed for detecting general SVGs but do not incorporate spatial features such as boundaries or centroids. To assess the concordance between the two *SpNeigh* approaches, we analyzed cluster 0 (meningeal fibroblasts) using both boundary- and centroid-derived spatial weights (Fig.3d). We then compared the ranks of genes based on Z1 coefficients from `RunSpatialDE` and normalized SEI values from `ComputeSpatialEnrichmentIndex`. The rank correlation was high, with Spearman coefficients of 0.75 (boundary weights) and 0.80 (centroid weights) (Fig. 3e), suggesting strong agreement between the two methods.

Notably, `RunSpatialDE` identified genes with increasing or decreasing trends along spatial gradients. For example, *Dcn* and *Col1a1*, markers of fibroblasts enriched near boundaries, were ranked among the top positive spatial DE genes, whereas *Slc17a7* and *Car4*, markers of cortical neurons, showed negative spatial trends (Fig. 3f). The same fibroblast markers were also recovered by `ComputeSpatialEnrichmentIndex` (Fig. 3g), reinforcing the biological relevance of both approaches.

Both functions are computationally efficient and completed in under one second per run. While `RunSpatialDE` provides statistical inference and directionality of expression changes, `ComputeSpatialEnrichmentIndex` offers a complementary enrichment-based perspective that may capture biologically relevant genes overlooked by hypothesis testing. Together, these methods offer a robust and flexible toolkit for modeling spatial expression gradients.

### 2.4 SpNeigh reveals immune microenvironment differences between breast tumor and DCIS neighborhoods

The tumor microenvironment plays a critical role in cancer progression by influencing tumor growth, immune surveillance, and mechanisms of immune evasion [18, 19]. Although most invasive breast cancers arise from ductal carcinoma in situ (DCIS), only 10–30% of DCIS cases eventually progress to invasive disease [20]. To demonstrate the utility of neighborhood analysis in *SpNeigh*, we applied it to a processed human breast cancer Xenium dataset with *SingleR*-based cell type annotations from Cheng et al. [21].

We focused on the spatial neighborhoods surrounding tumor and DCIS regions (Fig. 4a). Neighborhood ring regions were computed for tumor and DCIS separately, and cells located within these rings were extracted for downstream analysis (Fig. 4b). Cell type composition within each neighborhood was summarized (Fig. 4c). In the tumor neighborhood, fibroblasts were most abundant (29%), followed by T cells (14%), DCIS cells (13%), tumor cells (12%), and macrophages (11%). In the DCIS neighborhood, fibroblasts (34%) and T cells (17%) remained the top two cell types, with macrophages (11%) and DCIS cells (7%) also present.

**Fig. 4.**
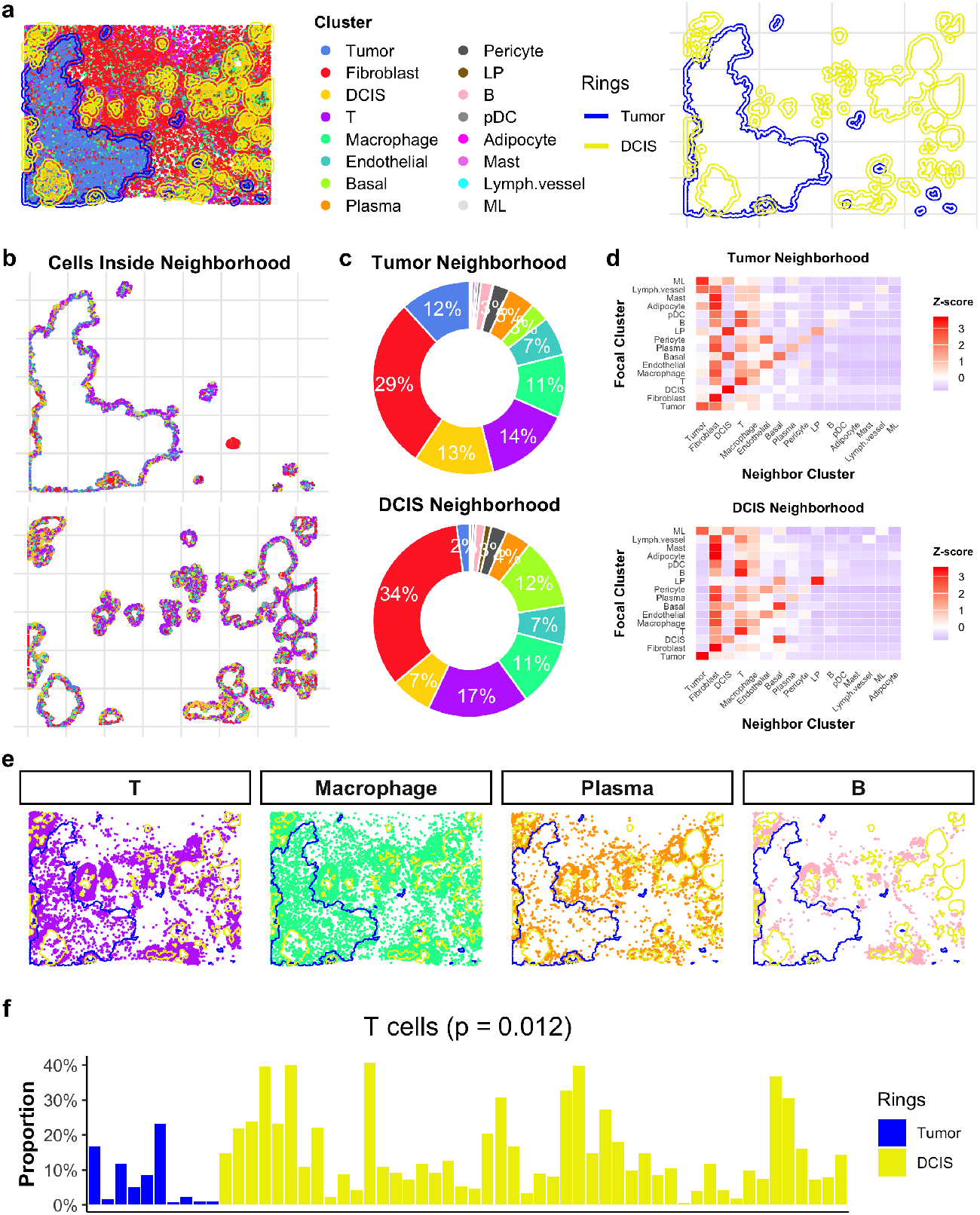
Neighborhood analysis reveals immune cell distribution differences around tumor and DCIS regions. **a**. Spatial plot of Xenium breast cancer tissue colored by cell type annotations. Neighbor ring regions surrounding tumor and ductal carcinoma in situ (DCIS) are shown separately for clarity. Blue indicates tumor-associated rings; yellow indicates DCIS-associated rings. **b**. Cells located within tumor and DCIS neighbor rings. **c**. Overall cell type proportions in tumor and DCIS neighbor rings. Cells from all rings are aggregated for each condition to compute relative frequencies (labels shown for proportions ≥2%). **d**. Row-scaled spatial interaction matrices showing the frequency of neighboring cell types around each focal cell type within tumor and DCIS neighborhoods. **e**. Spatial distribution of major immune cell types including T cells, macrophages, plasma cells, and B cells. Blue and yellow polygons indicate boundaries of tumor and DCIS, respectively. Notably, T, B, and plasma cells appear more enriched near DCIS than tumor regions. **f**. Bar plot comparing T cell proportions between tumor and DCIS neighbor rings. Blue bars represent tumor-associated neighborhoods and yellow bars represent DCIS. The T cell proportion is significantly higher near DCIS (Student’s t-test, p = 0.012).

To investigate cellular interactions within these neighborhoods, we computed spatial interaction matrices using `ComputeSpatialInteractionMatrix`. The row-scaled heatmap (Fig. 4d) revealed distinct interaction patterns: DCIS cells in the tumor neighborhood primarily interacted with other DCIS cells, whereas DCIS in the DCIS neighborhood showed interactions with basal cells, DCIS, and fibroblasts. These differences suggest potential heterogeneity in DCIS subpopulations depending on their proximity to tumor regions. Moreover, immune cells such as T, B, plasma, and macrophage populations were observed to interact with T cells in both neighborhoods, indicating potential immune modulation.

To assess immune infiltration visually, we plotted the spatial distributions of major immune cell types (T, B, plasma, and macrophages) relative to tumor and DCIS boundaries (Fig. 4e). Notably, T, B, and plasma cells appeared more enriched around DCIS regions. To quantify these differences, we compared cell type proportions using a two-sided Student’s t-test. T cells were significantly more abundant in DCIS neighborhoods than in tumor neighborhoods (p = 0.012; Fig. 4f), whereas no significant differences were observed for macrophages (p = 0.166), plasma cells (p = 0.413), or B cells (p = 0.448). These results suggest a potentially protective immune environment surrounding DCIS, which may contribute to its limited progression into invasive carcinoma.

Overall, this case study demonstrates how *SpNeigh* can reveal subtle yet biologically meaningful differences in the cellular and immune landscapes between tumor and pre-invasive lesions using neighborhood-aware spatial analysis.

### 2.5 SpNeigh reveals expression changes during early-stage tumor development from DCIS

To investigate transcriptional changes during the transition from DCIS to invasive breast cancer, we analyzed tumor and DCIS cells in the same human breast cancer Xenium dataset. Boundaries were identified for both tumor and DCIS populations, revealing one major tumor region and multiple spatially distinct DCIS regions (Fig. 5a). As only a subset of DCIS cases progress to invasive carcinoma, we hypothesized that DCIS cells near tumors might show distinct gene expression signatures.

**Fig. 5.**
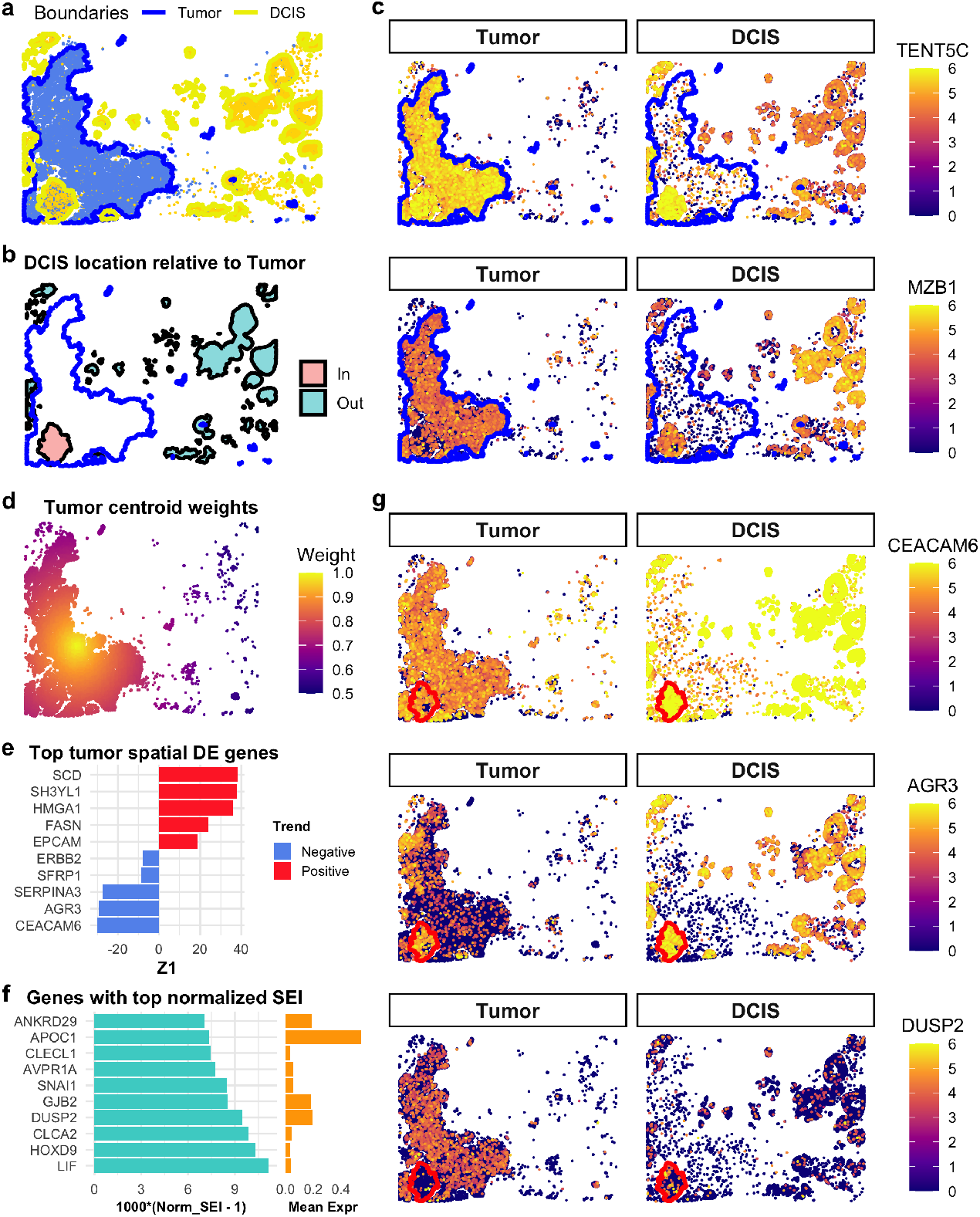
Spatial differential expression analysis of DCIS and tumor cells. **a**. Spatial plot of tumor and DCIS regions with boundaries. Blue and yellow polygons represent tumor and DCIS boundaries, respectively. **b**. Spatial mapping of DCIS boundary regions located inside versus outside tumor boundaries. The intratumoral DCIS region is highlighted in red; extratumoral regions are shown in green. **c**. Expression of TENT5C and MZB1 in DCIS cells located inside versus outside tumor boundaries. TENT5C is upregulated in DCIS cells within or near tumor regions, while MZB1 is downregulated. Expression profiles of these genes in intratumoral DCIS resemble those of tumor cells. **d**. Spatial weights based on distance to the centroid of tumor cells. Weights range from 0.5 (farther from the centroid) to 1 (closer to the centroid). **e**. Top 10 spatially varying genes in tumor cells ranked by Z1 from centroid-weighted RunSpatialDE. The sign of Z1 (x-axis) indicates the direction of spatial expression trend. **f**. Top 10 spatially enriched genes in tumor cells based on normalized SEI. To aid visualization, the x-axis shows 1000 times (normalized SEI - 1), emphasizing subtle variation near the baseline value of 1. **g**. Expression of CEACAM6, AGR3, and DUSP2 in tumor and DCIS cells. In tumor cells, CEACAM6 and AGR3 expression decreases with increasing distance from the centroid, while DUSP2 increases. CEACAM6 is also a DCIS marker gene and shows distinct expression between tumor and DCIS populations. The red polygon indicates the boundary of the intratumoral DCIS.

One DCIS region was located entirely within the tumor boundary (highlighted in red), while the remaining DCIS regions were outside the tumor region (green) (Fig. 5b). We performed differential expression analysis using `RunLimmaDE` to compare DCIS cells located inside versus outside the tumor boundary. Compared to DCIS cells distant from the tumor, tumor-adjacent DCIS cells showed elevated expression of *TENT5C* and reduced expression of *MZB1* (Fig. 5c). Notably, expression levels of these two genes in the tumor-adjacent DCIS cells resembled those in tumor cells, suggesting early transcriptional reprogramming. Previous work based on the same dataset also identified *MZB1* as a specific marker of DCIS regions distant from the tumor [1]. Interestingly, both *TENT5C* and *MZB1* are known plasma cell markers [22, 23] (Supplementary Fig. 2), and the observed shifts in their expression may reflect immune microenvironment remodeling associated with early tumor progression.

To further examine intratumoral heterogeneity, we focused on spatial gradients within the tumor itself. We reasoned that tumor cells near the boundary may represent earlier developmental stages, while cells closer to the tumor centroid may be more advanced. We therefore computed centroid-based spatial weights for tumor cells (Fig. 5d) and applied both `RunSpatialDE` and `ComputeSpatialEnrichmentIndex` to identify spatially varying and enriched genes along this inferred developmental axis.

*CEACAM6* and *AGR3* were among the top genes showing decreasing expression along the centroid weights (Fig. 5e), while *DUSP2* emerged as a highly enriched gene in cells near the centroid with high average expression across tumor cells (Fig. 5f). Visualization of these genes in both DCIS and tumor cells showed distinct spatial expression patterns (Fig. 5g). Notably, the DCIS region located inside the tumor boundary contained both tumor and DCIS cells. *CEACAM6*, a canonical DCIS marker [1], was highly expressed in all DCIS cells, including those within the tumor region, but was downregulated in neighboring tumor cells, showing a clear negative trend (Z1 *<*0) along the centroid weights. *AGR3* expression also decreased progressively from DCIS cells to adjacent tumor cells and was lowest in tumor cells near the centroid. In contrast, *DUSP2* was most highly expressed in centroid-proximal tumor cells and nearly absent in DCIS cells.

These results highlight the ability of *SpNeigh* to reveal spatial expression gradients and transitional states during tumor evolution. By modeling spatial context and neighborhood relationships, *SpNeigh* provides valuable insights into early tumor development and cellular reprogramming at the invasive front.

### 2.6 SpNeigh reveals gene expression changes along spatial gradients of liver zonation

The liver exhibits a highly structured lobular architecture, consisting of three zones: periportal (zone 1), midlobular (zone 2), and pericentral (zone 3) [24]. Midlobular hepatocytes are thought to contribute to hepatocyte renewal during homeostasis and regeneration [25]. To investigate zonation-associated gene expression patterns, we applied the spatial differential expression analysis in *SpNeigh* to a publicly available MERFISH dataset of healthy human liver tissue [26].

Low-quality cells were filtered out, and 23,144 hepatocytes were retained for downstream analysis. The hepatocyte subpopulations annotated by the original authors—hepatocyte 1, 2, and 3—correspond to periportal, midlobular, and pericentral zones, respectively. Boundaries for each hepatocyte population were computed and visualized in spatial plots (Fig.6a). We then computed spatial weights to the boundaries of each population (Fig.6b), and applied `RunSpatialDE` to identify spatially varying genes along these gradients.

The top 10 spatially differential genes for each hepatocyte population, identified using boundary-based weights, showed clear zonation-related expression trends (Supplementary Fig. 3a). For example, *ASS1* and *SDS* were enriched near the boundaries of hepatocyte 1, with highest expression in hepatocyte 1 and intermediate expression in hepatocyte 2. Similarly, *CYP1A1* and *CYP3A4* were enriched near the boundaries of hepatocytes 2 and 3, with highest expression in hepatocyte 3 and intermediate levels in hepatocyte 2 (Fig. 6c, Supplementary Fig. 3a).

**Fig. 6.**
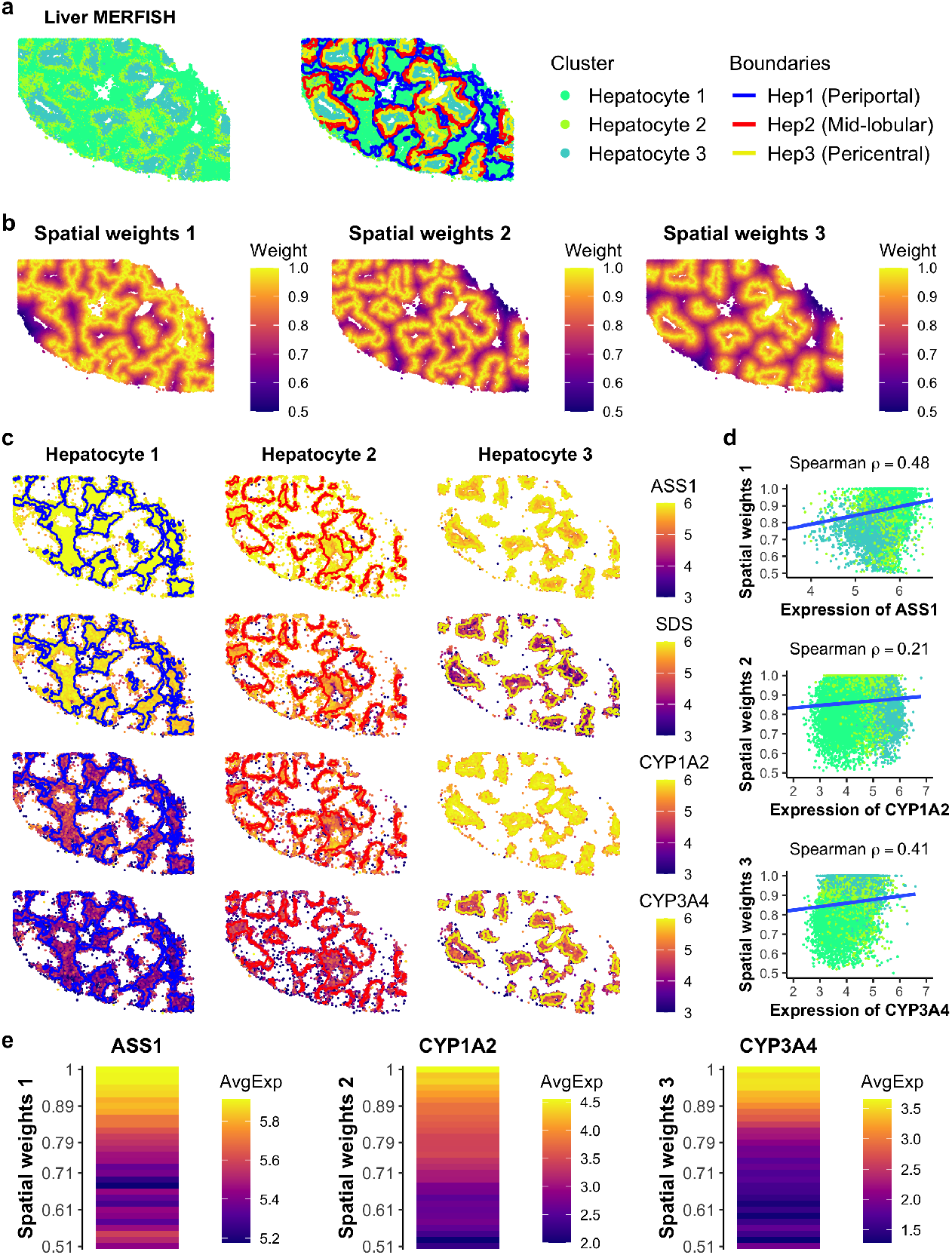
Spatial differential expression analysis of hepatocytes in healthy liver using MERFISH. **a**. Spatial plot of hepatocyte subpopulations and their detected boundaries in a publicly available MERFISH liver dataset. Hepatocyte 1, 2, and 3 correspond to periportal, mid-lobular, and pericentral zones, respectively. Blue, red, and yellow lines indicate the boundaries of hepatocyte populations 1, 2, and 3, respectively. **b**. Spatial weights derived from distances to the boundaries of each hepatocyte zone. Higher values indicate proximity to the boundary; a weight of 1 denotes a cell located precisely on the boundary. **c**. Spatial expression of top-ranked spatially varying genes along boundary-based weights. ASS1 and SDS are enriched in periportal hepatocytes (zone 1); CYP1A1 and CYP3A4 are enriched in mid-lobular and pericentral hepatocytes (zones 2 and 3). **d**. Scatter plots showing normalized expression of marker genes versus spatial weights. Spearman correlation coefficients quantify the association between expression and distance. Only cells with nonzero expression are shown. **e**. Heatmap of average gene expression across spatial weight bins. ASS1, CYP1A1, and CYP3A4 illustrate representative spatial expression patterns along boundary weights of zones 1, 2, and 3, respectively.

Expression levels of representative genes were correlated with their corresponding spatial weights using Spearman correlation. The correlations for *ASS1, CYP1A1*, and *CYP3A4* with boundary weights for hepatocytes 1, 2, and 3 were 0.48, 0.21, and 0.41, respectively (Fig.6d). Binned expression plots further demonstrated smooth gradients in gene expression along these spatial axes (Fig.6e). Additional examples include *GLS2*, which showed peak expression in hepatocyte 1 and intermediate expression in hepatocyte 2, and *CYP2E1*, which showed the opposite trend, peaking in hepatocyte 3 (Supplementary Fig. 3b).

These findings are consistent with previously reported liver zonation markers, including periportal genes (*GLS2, ASS1, SDS*) and pericentral genes (*CYP1A1, CYP3A4, CYP2E1*) [26–30]. Interestingly, although canonical midlobular markers such as *HAMP, HAMP2*, and *CCND1* [25, 27] were not included in the 317-gene MERFISH panel, we identified *COL1A1* as a potential negative marker for hepatocyte 2 (midlobular zone). Its expression was lowest in hepatocyte 2 and showed a negative trend along the spatial weights to hepatocyte 2 boundaries (Supplementary Fig. 3a,b). As *COL1A1* is a known marker of hepatic stellate cells—cell types implicated in regulating liver zonation [31]—its expression in hepatocytes may reflect zone-specific regulatory interactions.

Together, these results demonstrate the ability of *SpNeigh* to reveal spatial expression gradients aligned with liver zonation and highlight its utility in boundary-aware analysis of spatial transcriptomics data.

## 3 Discussion

Spatial transcriptomics technologies have transformed our ability to study gene expression in tissue architecture, but extracting biological meaning from spatial organization remains methodologically challenging. *SpNeigh* introduces a flexible and interpretable framework for spatial neighborhood modeling, integrating local boundary detection, distance-aware weighting, spatial differential expression, and enrichment analysis. Through modular components, *SpNeigh* enables the investigation of cell-cell interactions, microenvironmental heterogeneity, and spatially varying transcriptional programs across diverse tissue types and spatial platforms.

Our analysis demonstrates that *SpNeigh* effectively captures biologically relevant spatial features across a range of datasets. In mouse brain, boundary-aware modeling identified intermediate populations and layered expression gradients near white matter boundaries. In human breast cancer, neighborhood ring analysis revealed distinct immune microenvironments surrounding tumor and DCIS regions, highlighting spatially structured immune infiltration. In healthy liver, *SpNeigh* recovered known zonation patterns and identified new spatially varying genes, including zone-specific markers and potential regulatory candidates. Together, these examples showcase the utility of *SpNeigh* for discovering fine-grained transcriptional variation driven by spatial context.

Compared to existing methods such as *SpatialDE* [6], *SPARK* [7], and *Giotto* [4], which primarily focus on global spatial variability or spatial autocorrelation, *SpNeigh* is uniquely designed for neighborhood-aware modeling and boundary-centric analysis. The incorporation of spatial weights anchored to biologically meaningful structures—such as cluster boundaries or centroids—allows for directional testing of spatial expression gradients. Unlike Gaussian process-based methods that may suffer from scalability issues on large datasets, our use of spline-based linear modeling and weighted means ensures rapid computation without sacrificing interpretability.

While *SpNeigh* introduces novel capabilities, several directions for future development remain. For example, automated region selection, dynamic modeling of cell-cell interactions, and multivariate spatial analyses could further enhance its applicability. The current framework focuses primarily on 2D spatial datasets, but extensions to 3D tissue volumes and time-series spatial data are conceptually straightforward. Additionally, integrating histological features or multi-modal data may enable richer modeling of spatial niches.

In summary, *SpNeigh* addresses a critical need for interpretable and efficient spatial modeling tools that account for local context and region-specific variation. Its flexible design and compatibility with modern spatial platforms make it a valuable addition to the spatial transcriptomics toolkit, supporting a broad range of biological and clinical investigations.

## 4 Methods

### 4.1 SpNeigh framework for spatial neighborhood modeling

#### 4.1.1 Spatial boundary detection and outlier removal

To identify the spatial boundaries of a cell population or cluster, we first remove spatial outliers based on local density. Let **s**_*c*_ denote the spatial coordinate of cell *c*. For each cell, we compute the average Euclidean distance to its *k* nearest neighbors using the *k*-nearest neighbor (k-NN) algorithm:

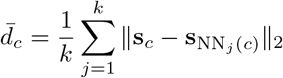

where NN_*j*_(*c*) denotes the *j*-th nearest neighbor of cell *c*, and 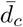is the mean k-NN distance. Cells with 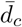 greater than a user-defined threshold *τ* are considered spatial outliers and removed.

After filtering, spatial boundaries are identified using the concave hull algorithm [32], which estimates a smoothed outer boundary encompassing each connected spatial region. If a cluster contains multiple disjoint regions, subregions can be detected using either DBSCAN [33] (density-based clustering) or *k*-means. Each subregion is then processed independently to construct its boundary polygon.

The resulting boundaries are represented as a set of spatial polygons or LINESTRINGs, which are used for computing spatial weights, enrichment indices, or visualization.

#### 4.1.2 Spatial neighborhood interaction matrix

To characterize spatial relationships between distinct cell clusters, we compute a neighborhood interaction matrix based on k-NN counts. For each cell, we identify its *k* nearest spatial neighbors using Euclidean distance and record their cluster identities.

Let 𝒞 denote the set of all clusters, and define 𝒞_*i*_ as the set of all cells belonging to cluster *I* ∈ 𝒞. Let N_*k*_(*c*) be the set of the *k* nearest neighbors of cell *c*, and let cluster(*n*) denote the cluster identity of neighbor cell *n*.

We define the spatial interaction matrix *M* ∈ ℝ^|*𝒞*|*×*|*𝒞*|^ such that:

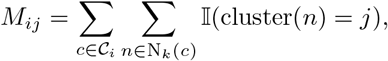

where 𝕀 (·) is the indicator function, equal to 1 when the condition is true and 0 otherwise.

This matrix *M* quantifies how often cells from cluster *i* are spatially adjacent to cells from cluster *j* in their local neighborhood. It can be visualized directly or row-normalized (e.g., Z-scored) to highlight enriched spatial associations.

### 4.1.3 Spatial weights from boundaries and centroids

To model gene expression variation relative to spatial structures, we assign each cell a spatial weight based on its distance to a reference geometry such as a boundary or a centroid. Let **s**_*c*_ denote the 2D spatial coordinate of cell *c*.

For centroid-based weighting, we compute the centroid position of a selected cell population as:

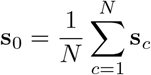

where *N* is the number of selected cells. The Euclidean distance between each cell and the centroid is:

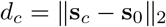

For boundary-based weighting, the shortest distance from cell *c* to the nearest boundary location is:

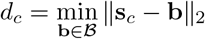

where ℬ denotes the set of all boundary points, and **b** ∈ ℬ represents a specific boundary location.

To ensure comparability across regions of varying scale, we apply min-max normalization:

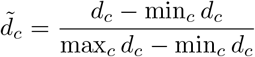

The spatial weight *w*_*c*_ for cell *c* is then computed using one of several decay functions applied to the normalized distance 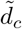:

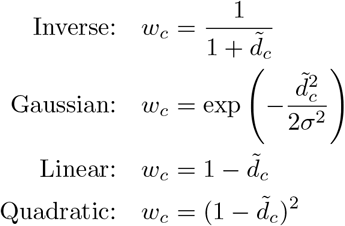

where *σ* is a user-defined scale parameter used in the Gaussian kernel. These weights can be used directly in enrichment analysis or as input covariates in downstream modeling of gene expression.

#### 4.1.4 Gradient-based spatial differential expression

To identify genes with expression patterns that vary smoothly along a spatial distance gradient, we modeled gene expression as a continuous function of spatial distance using a reparameterized spline basis, similar to approaches used in pseudotime trajectory analysis [34]. Let **t** = (*t*_1_, *t*_2_, …, *t*_*n*_) be a vector of spatial distances for *n* cells, such as proximity to a boundary or centroid. We construct a natural cubic spline basis ns(**t**) using the `splines::ns()` function with *k* degrees of freedom (default *k* = 3), yielding a design matrix *X* ∈ ℝ^*n×k*^ .

An augmented matrix *A* = [**1, t**, *X*] is formed by concatenating an intercept vector **1**, the distance vector **t**, and the spline basis. We then apply QR decomposition to orthonormalize the matrix (excluding the intercept), producing a final design matrix *Z* ∈ ℝ^*n×k*^, with orthogonal and unit-norm columns. The first column of *Z* is adjusted to be positively correlated with **t**, ensuring alignment with the primary gradient direction.

For each gene *g*, expression is modeled as a linear combination:

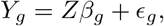

where *Y*_*g*_ ∈ ℝ^*n*^ is the expression vector for gene *g, β*_*g*_ ∈ ℝ^*k*^ is the coefficient vector, and *ϵ*_*g*_ represents residual errors. Hypothesis testing is performed across all coefficients using the *limma* [8] framework.

To aid interpretability, we report the sign of the first coefficient *β*_*g*,1_ as the primary trend direction. A positive value indicates increasing expression with distance, while a negative value indicates decreasing expression.

#### 4.1.5 Spatial enrichment index

To quantify the spatial bias of a gene toward spatially defined regions (e.g., near boundaries or centroids), we compute a SEI for each gene. For gene *g* and cell *c*, let *y*_*gc*_ denote the expression level of gene *g* in cell *c*, and let *w*_*c*_ denote the spatial weight assigned to cell *c*. The SEI is defined as the weighted mean expression of gene *g* across all cells:

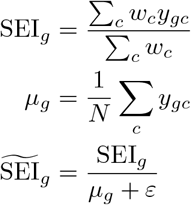

where *µ*_*g*_ is the unweighted mean expression of gene *g, N* is the number of cells, and *ε* is a small constant (e.g., 10^−6^) added to avoid division by zero. The normalized SEI, 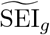, is used to compare spatial enrichment independently of absolute gene expression levels. Genes with large normalized SEI values are considered spatially enriched relative to the provided spatial weights.

### 4.2 Spatial transcriptomics and single-cell datasets

The following spatial transcriptomics and scRNA-seq datasets have been used in this study:

1. The raw mouse brain tiny Xenium dataset was downloaded from 10x Genomics dataset website: https://www.10xgenomics.com/datasets/fresh-frozen-mouse-brain-for-xenium-explorer-demo-1-standard.
2. The processed human breast cancer Xenium dataset (Sample 1) was obtained from Cheng et al [21], and the raw data was from 10x Genomics dataset website: https://www.10xgenomics.com/products/xenium-in-situ/preview-dataset-human-breast [1].
3. The processed human healthy liver MERFISH dataset was downloaded from: https://datadryad.org/stash/dataset/doi:10.5061/dryad.37pvmcvsg [26], and the hepatocytes were used in this study.
4. The processed mouse brain scRNA-seq dataset was downloaded from Gene Expression Omnibus with accession number of GSE185862 [14]. A subset of 10,000 cells was extracted from the processed scRNA-seq data as a reference for cell type annotation of mouse brain Xenium data.

### 4.3 Data preparation for SpNeigh

The mouse brain tiny Xenium dataset was processed using the *Seurat* (v5.1.0) pipeline [12]. Cells with fewer than 10 detected genes or fewer than 15 total counts were excluded as part of the quality control procedure. The filtered dataset was normalized using `NormalizeData`, followed by variance stabilization with `ScaleData`. Dimensionality reduction was performed via `RunPCA` and `RunUMAP`, and clustering was carried out using `FindNeighbors` and `FindClusters`.

To perform cell type annotation at the single-cell level, a small reference scRNA-seq dataset consisting of 10,000 cells was sampled from the mouse brain dataset published by Yao et al. [14], which contains approximately 1 million cells. During the sampling process, at least 50 cells were retained for each cell type to ensure sufficient representation of all annotated populations in the reference. Annotation of the Xenium dataset was then conducted using *SingleR* (v2.8.0) [13], and predicted cell labels were manually merged into broader categories following the taxonomy defined by Yao et al. to reduce annotation granularity.

The processed human breast cancer Xenium dataset from Cheng et al., including *SingleR*-based cell type annotations, was used directly for downstream analysis.

The human healthy liver MERFISH dataset underwent similar quality control, removing cells with low gene or UMI counts. After normalization with `NormalizeData`, author-annotated hepatocyte populations were extracted for downstream analysis.

For all datasets, the spatial coordinate data frame (with columns `x`, `y`, `cell`, and `cluster`) was extracted from the *Seurat* object using the `ExtractCoords` function from *SpNeigh*. The normalized expression matrix was retrieved using `GetAssayData` from *Seurat*. These coordinate and expression inputs were used throughout the *SpNeigh* analysis pipeline.

### 4.4 Analysis of mouse brain Xenium dataset using SpNeigh

To identify spatial boundaries for cluster 2 in the mouse brain Xenium dataset, we applied the `GetBoundary` function with a refined `eps` parameter of 120 to detect spatially coherent outlier-free contours. Boundary polygons were subsequently constructed from these points using the `BuildBoundaryPoly` function. To define the surrounding neighborhood, ring regions were generated using the `GetRingRegion` function, which by default applies a 100-unit buffer to the original boundary polygons via `GetOuterBoundary`.

Cells located within the cluster 2 boundaries or neighborhood rings were extracted using the `GetCellsInside` function. Cell composition statistics for these regions were computed with `StatsCellsInside`. Differential expression analysis between cluster 2 cells located inside the boundaries and those in the surrounding rings was conducted using `RunLimmaDE`.

To further dissect spatial structure, the primary boundary polygon of cluster 2 was segmented into individual edges using the `SplitBoundaryPolyByAnchor` function. Focusing on the cerebral cortex, spatial weights to a specific edge (edge 2) were computed for relevant cells using `ComputeBoundaryWeights`. Finally, spatial differential expression along this boundary gradient was assessed using the `RunSpatialDE` function.

### 4.5 Benchmarking differential expression and spatial modeling in SpNeigh

To evaluate the performance of differential expression analysis in *SpNeigh*, we compared the `RunLimmaDE` function against the widely used `FindAllMarkers` function from the *Seurat* package. Both methods were applied to the mouse brain Xenium dataset using consistent parameter settings (`min.pct = 0`) to identify marker genes for each cell type based on cluster-level annotations. We recorded the total runtime for each method and computed the Spearman correlation between log_2_ fold changes (log_2_FC) across all identified marker genes to assess the concordance between the two approaches.

The spatial modeling functions `RunSpatialDE` and `ComputeSpatialEnrichmentIndex` represent novel contributions introduced in *SpNeigh*, and are not directly comparable to existing methods. While *SpatialDE* [6] is a popular tool for identifying SVGs, it is primarily designed for unsupervised detection of broad spatial patterns and does not explicitly account for spatial boundaries or region-specific structures. Moreover, *SpatialDE* ‘s reliance on Gaussian process modeling introduces substantial computational overhead, limiting its scalability for large-scale datasets such as Xenium or MERFISH. In contrast, *SpNeigh* models spatial gradients directly using user-defined spatial weights derived from boundaries or centroids, enabling region-aware and computationally efficient analysis.

We assessed the concordance between `RunSpatialDE` and `ComputeSpatialEnrichmentIndex` by analyzing cluster 0 cells from the mouse brain Xenium dataset. Spatial weights were computed based on both boundary and centroid distances. Genes were ranked by Z_1_ coefficients from `RunSpatialDE` and by normalized SEI from `ComputeSpatialEnrichmentIndex`. Spearman correlation was used to quantify agreement between rankings. Additionally, top-ranked genes were manually inspected to confirm spatial enrichment patterns and biological relevance. As `RunSpatialDE` shares the same *limma*-based modeling framework as `RunLimmaDE`, its runtime was not reported separately due to similar computational efficiency.

### 4.6 Analysis of human breast cancer Xenium dataset using SpNeigh

#### 4.6.1 Neighborhood analysis of tumor and DCIS regions

To characterize the local neighborhoods of tumor and DCIS in the human breast cancer dataset, boundaries were computed for all spatially distinct tumor and DCIS regions using the `GetBoundary` function. Neighborhood ring regions surrounding the tumor and DCIS were then generated using `GetRingRegion`, which constructs outer boundaries at a specified distance from the original boundaries. Cells located within these ring regions were extracted with `GetCellsInside`, and their cell type composition was quantified using `StatsCellsInside`. Donut plots were generated with `PlotStatsPie` (`plot donut = TRUE`) to visualize the proportion of each cell type in the tumor and DCIS neighborhoods. Additionally, Student’s t-test was used to assess statistically significant differences in the proportion of each cell type between the tumor and DCIS ring regions.

To quantify cell–cell interactions within these local neighborhoods, the `ComputeSpatialInteractionMatrix` function was applied to the ring-localized cells, generating a matrix of interactions between focal clusters and their neighboring clusters. This matrix was row-scaled and visualized using `PlotInteractionMatrix`.

#### 4.6.2 Differential expression and spatial modeling of tumor and DCIS cells

To explore molecular differences between DCIS cells in distinct spatial locations, differential expression analysis was performed using `RunLimmaDE` to compare DCIS cells located inside versus outside of tumor boundaries.

To further investigate intra-tumoral heterogeneity and spatial gradients in gene expression, `ComputeCentroidWeights` was used to compute spatial weights based on proximity to the tumor centroid. These weights were used as a covariate in `RunSpatialDE` to identify genes with spatially varying expression patterns along the centroid-based gradient. Differentially expressed genes were ranked by adjusted p-values. In parallel, `ComputeSpatialEnrichmentIndex` was applied to the same cells using centroid weights to identify spatially enriched genes, which were ranked based on normalized SEI scores.

### 4.7 Analysis of human healthy liver MERFISH dataset using SpNeigh

To investigate spatially varying gene expression along liver zonation gradients, boundaries were first identified for hepatocyte populations corresponding to the periportal, mid-lobular, and pericentral zones using the `GetBoundary` function. Spatial weights reflecting proximity to each zone-specific boundary were then computed with `ComputeBoundaryWeights`. These weights were used as input to `RunSpatialDE`, which models gene expression as a smooth function of spatial distance using natural splines. To assess the strength and direction of spatial variation, Spearman correlation coefficients were calculated between normalized gene expression and spatial weights for the top-ranked genes identified by `RunSpatialDE`. Finally, `PlotSpatialExpression` was used to visualize the average expression trends of selected genes along the spatial gradients.

## Acknowledgements

We acknowledge the support of ChatGPT-4o (OpenAI) for assistance with code development, documentation, and language refinement of this manuscript.

## Data availability

In this study, we used publicly available datasets as detailed in Methods. Mouse brain tiny Xenium dataset: https://www.10xgenomics.com/datasets/fresh-frozen-mouse-brain-for-xenium-explorer-demo-1-standard. Human breast cancer Xenium dataset: https://www.10xgenomics.com/products/xenium-in-situ/preview-dataset-human-breast [1]. Cheng et al annotation for human breast cancer Xenium dataset: https://github.com/jinming-cheng/cheng_annotation_bmc_bioinfo [21]. Human healthy liver MERFISH dataset: https://datadryad.org/stash/dataset/doi:10.5061/dryad.37pvmcvsg [26]. Mouse brain scRNA-seq dataset: GSE185862 [14].

## Code availability

The SpNeigh package is available at this GitHub repository: https://github.com/jinming-cheng/SpNeigh. It is also available on Zenodo https://doi.org/10.5281/zenodo.17505217 [35]: . Source code for reproducing all figures and analyses in this manuscript is publicly available at: https://github.com/jinming-cheng/Rcodes for figures SpNeigh ms.

## Author contributions

J.C. conceived the SpNeigh package, conducted all the data analyses, generated all the figures and wrote the manuscript. P.K.H.C and N.L. discussed and supervised this study. All authors read and approved the final manuscript.

## 6 Supplementary

**Supplementary Table. 1.** Manually merged SingleR cell type annotation for mouse brain.

| New_Merged_CellType | Raw_SingleR_CellType |
| --- | --- |
| Hippocampus | CA1-ProS, CA2-IG-FC, CA3, DG, SUB-ProS, CT SUB |
| L2/3 | L2 IT ENTl, L2 IT ENTm, L2/3 IT ENTl, L3 IT ENT, L2/3 IT CTX, L2/3 IT PPP, L2/3 IT RHP |
| L4/5/6 | L4 RSP-ACA, L4/5 IT CTX, L5 IT CTX, L5 PPP, L5 PT CTX, NP PPP, NP SUB, L6 CT CTX, L6 IT CTX, L6 IT ENTl, L5/6 IT TPE-ENT, L5/6 NP CTX, Car3 |
| L6b | L6b CTX, L6b/CT ENT |
| Astrocytes | Astro |
| Oligodendrocytes | Oligo |
| CGE | Lamp5, Sncg, Vip |
| MGE | Sst, Sst Chodl, Pvalb |
| Vascular | Endo, VLMC, SMC-Peri |
| Other | Meis2, CR, Micro-PVM |

**Supplementary Fig. 1.**
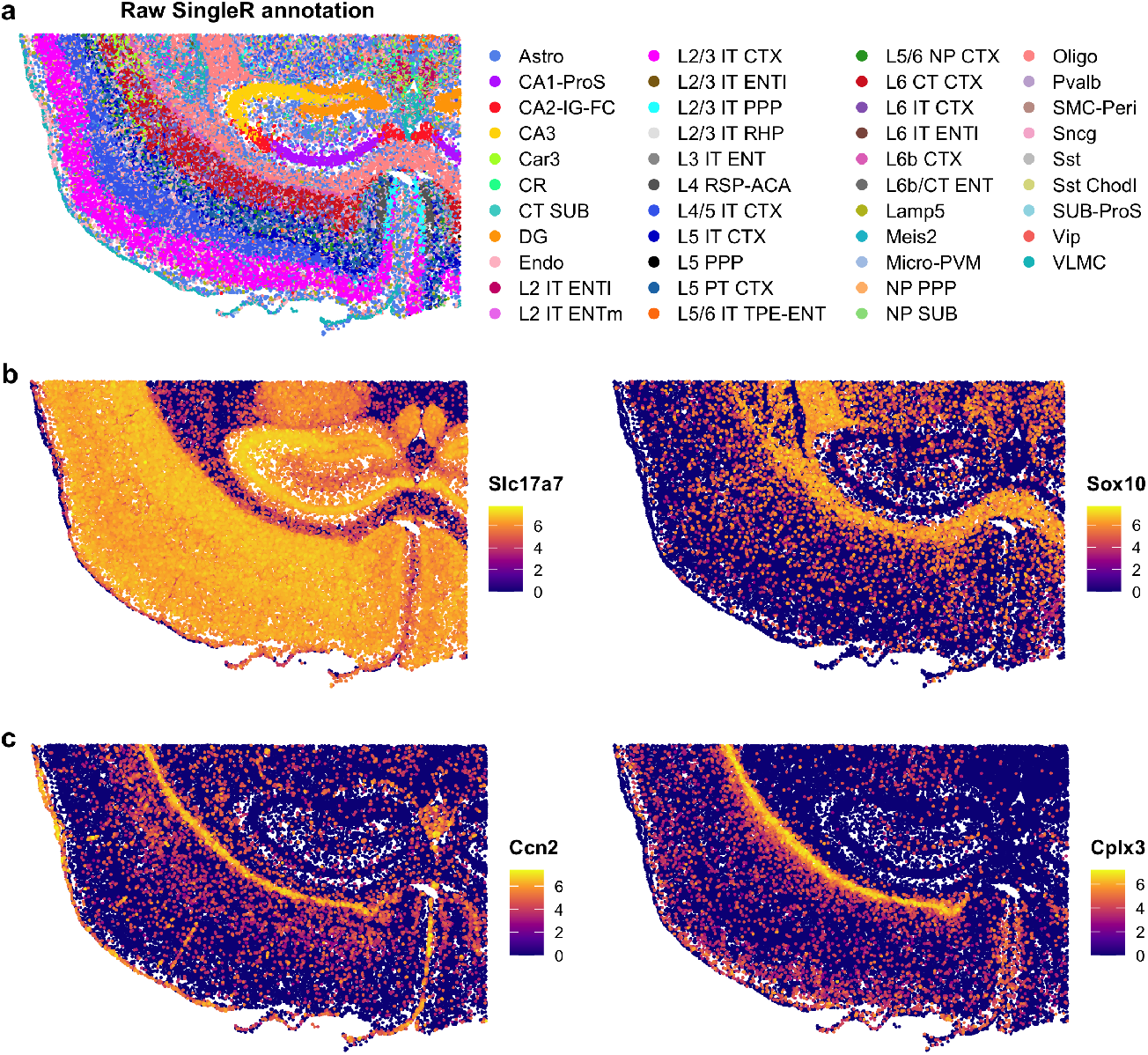
Cell type annotation and gene expression in mouse brain Xenium data. **a**. Spatial plot colored by cell-level annotations assigned using SingleR. **b**. Spatial expression of Slc17a7 and Sox10. Slc17a7 is a marker for excitatory neurons; Sox10 marks oligodendrocytes. **c**. Spatial expression of Ccn2 and Cplx3, genes enriched in boundary-associated neuronal populations.

**Supplementary Fig. 2.**
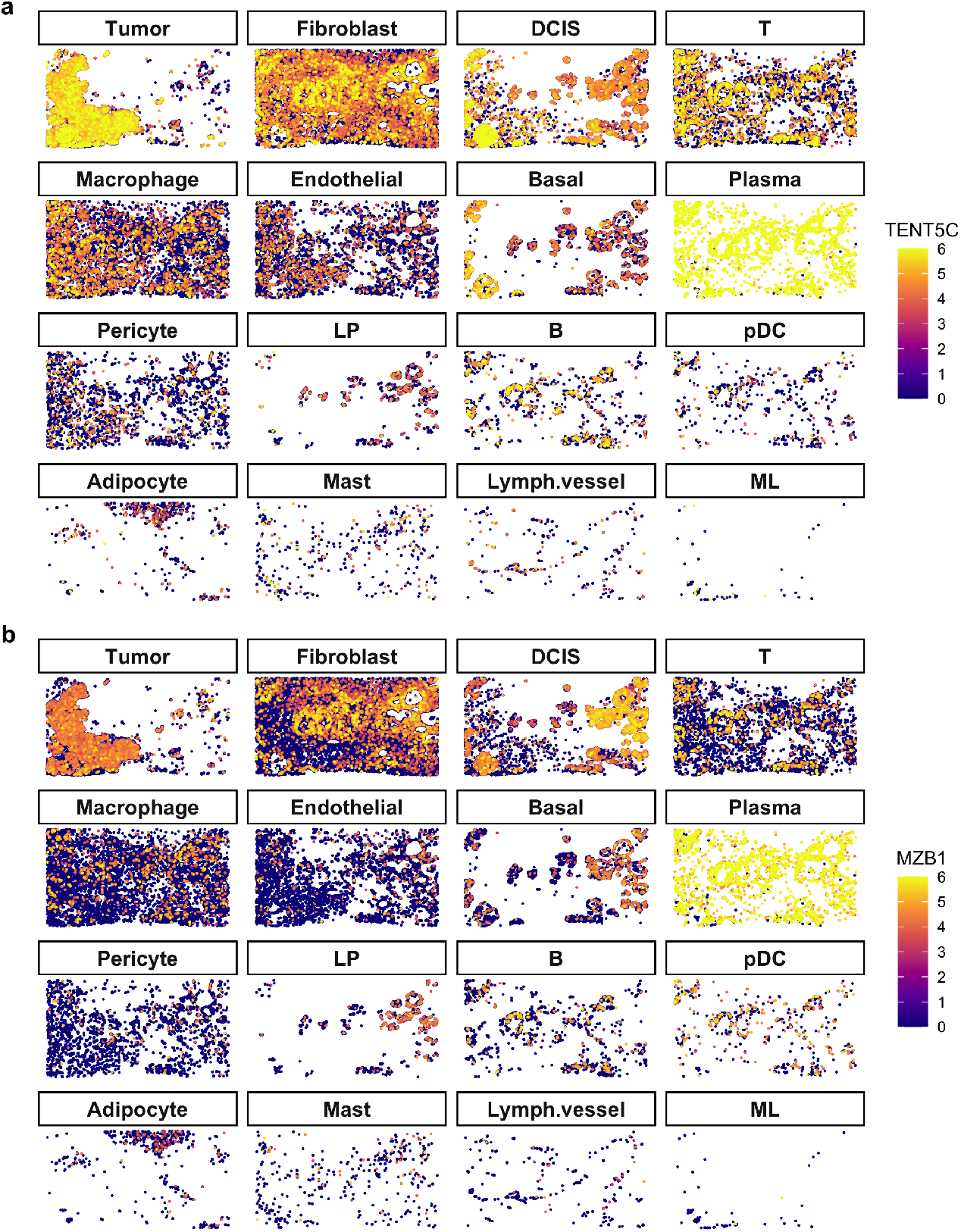
Expression of TENT5C and MZB1 across cell types in breast cancer Xenium data. **a**. Spatial expression of TENT5C across annotated cell types. **b**. Spatial expression of MZB1 across annotated cell types.

**Supplementary Fig. 3.**
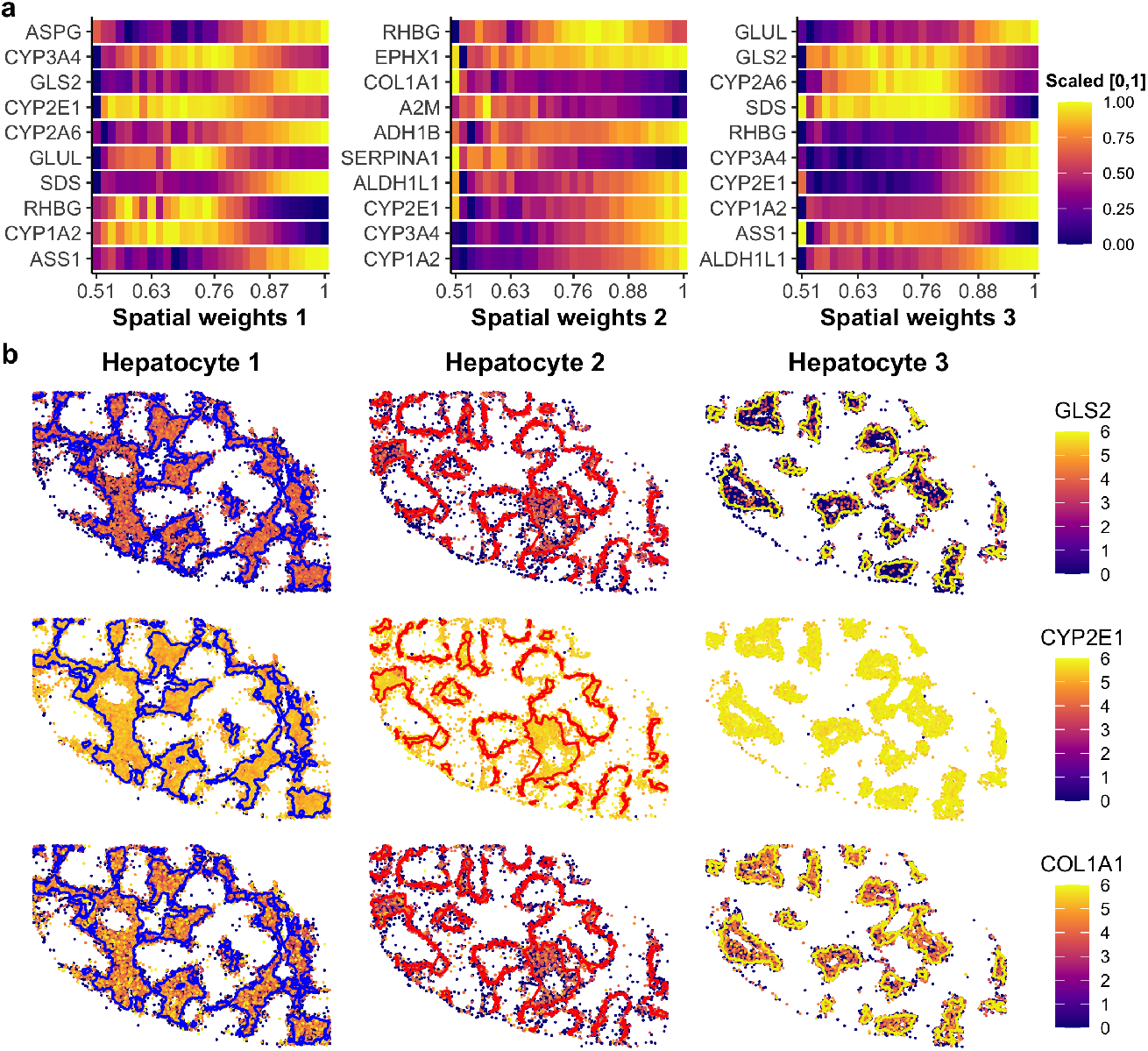
Spatial expression patterns of top spatially varying genes in liver hepatocytes. **a**. Heatmap showing the expression of the top 10 spatially differential genes along binned spatial weights for each hepatocyte population. Average expression values were rescaled to the range [0,1] to aid visualization. **b**. Spatial expression plots of GLS2, CYP2E1, and COL1A1 across the three hepatocyte populations. GLS2 is a periportal marker (hepatocyte 1), and CYP2E1 is a pericentral marker (hepatocyte 3); both show intermediate expression in mid-lobular hepatocytes (hepatocyte 2). COL1A1 shows lowest expression in hepatocyte 2. Blue, red, and yellow lines indicate the boundaries of hepatocyte populations 1, 2, and 3, respectively.

## References

[1] Janesick, A. et al. High resolution mapping of the tumor microenvironment using integrated single-cell, spatial and in situ analysis. Nat Commun 14, 8353 (2023).

[2] Moffitt, J. R. et al. High-throughput single-cell gene-expression profiling with multiplexed error-robust fluorescence in situ hybridization. Proc Natl Acad Sci U S A 113, 11046–51 (2016).

[3] Nagendran, M. et al. 1457 visium hd enables spatially resolved, single-cell scale resolution mapping of ffpe human breast cancer tissue. J Immunother Cancer 11, A1620 (2023).

[4] Dries, R. et al. Giotto: a toolbox for integrative analysis and visualization of spatial expression data. Genome Biol 22, 78 (2021).

[5] Palla, G. et al. Squidpy: a scalable framework for spatial omics analysis. Nat Methods 19, 171–178 (2022).

[6] Svensson, V., Teichmann, S. A. & Stegle, O. SpatialDE: identification of spatially variable genes. Nat Methods 15, 343–346 (2018).

[7] Sun, S., Zhu, J. & Zhou, X. Statistical analysis of spatial expression patterns for spatially resolved transcriptomic studies. Nat Methods 17, 193–200 (2020).

[8] Ritchie, M. E. et al. limma powers differential expression analyses for rna-sequencing and microarray studies. Nucleic Acids Res 43, e47 (2015).

[9] Yao, Z. et al. A high-resolution transcriptomic and spatial atlas of cell types in the whole mouse brain. Nature 624, 317–332 (2023).

[10] Shi, H. et al. Spatial atlas of the mouse central nervous system at molecular resolution. Nature 622, 552–561 (2023).

[11] Chen, A. et al. Spatiotemporal transcriptomic atlas of mouse organogenesis using DNA nanoball-patterned arrays. Cell 185, 1777–1792 e21 (2022).

[12] Hao, Y. et al. Dictionary learning for integrative, multimodal and scalable single-cell analysis. Nat Biotechnol 42, 293–304 (2024).

[13] Aran, D. et al. Reference-based analysis of lung single-cell sequencing reveals a transitional profibrotic macrophage. Nat Immunol 20, 163–172 (2019).

[14] Yao, Z. et al. A taxonomy of transcriptomic cell types across the isocortex and hippocampal formation. Cell 184, 3222–3241 e26 (2021).

[15] Qian, X. et al. Spatial transcriptomics reveals human cortical layer and area specification. Nature (2025).

[16] Viswanathan, S., Sheikh, A., Looger, L. L. & Kanold, P. O. Molecularly defined subplate neurons project both to thalamocortical recipient layers and thalamus. Cereb Cortex 27, 4759–4768 (2017).

[17] Kim, S. J. et al. A consensus definition for deep layer 6 excitatory neurons in mouse somatosensory, visual, and motor cortex. Cell Rep 44, 116167 (2025).

[18] Hanahan, D. & Weinberg, R. A. Hallmarks of cancer: the next generation. Cell 144, 646–74 (2011).

[19] Quail, D. F. & Joyce, J. A. Microenvironmental regulation of tumor progression and metastasis. Nat Med 19, 1423–37 (2013).

[20] Nolan, E., Lindeman, G. J. & Visvader, J. E. Deciphering breast cancer: from biology to the clinic. Cell 186, 1708–1728 (2023).

[21] Cheng, J., Jin, X., Smyth, G. K. & Chen, Y. Benchmarking cell type annotation methods for 10x Xenium spatial transcriptomics data. BMC Bioinformatics 26, 22 (2025).

[22] Bilska, A. et al. Immunoglobulin expression and the humoral immune response is regulated by the non-canonical poly(A) polymerase tent5c. Nat Commun 11, 2032 (2020).

[23] Andreani, V. et al. Cochaperone Mzb1 is a key effector of Blimp1 in plasma cell differentiation and beta1-integrin function. Proc Natl Acad Sci U S A 115, E9630–E9639 (2018).

[24] Martini, T., Naef, F. & Tchorz, J. S. Spatiotemporal metabolic liver zonation and consequences on pathophysiology. Annu Rev Pathol 18, 439–466 (2023).

[25] Wei, Y. et al. Liver homeostasis is maintained by midlobular zone 2 hepatocytes. Science 371 (2021).

[26] Watson, B. R. et al. Spatial transcriptomics of healthy and fibrotic human liver at single-cell resolution. Nat Commun 16, 319 (2025).

[27] Halpern, K. B. et al. Single-cell spatial reconstruction reveals global division of labour in the mammalian liver. Nature 542, 352–356 (2017).

[28] Xu, J. et al. A spatiotemporal atlas of mouse liver homeostasis and regeneration. Nat Genet 56, 953–969 (2024).

[29] Hildebrandt, F. et al. Spatial transcriptomics to define transcriptional patterns of zonation and structural components in the mouse liver. Nat Commun 12, 7046 (2021).

[30] Ang, C. H. et al. Self-maintenance of zonal hepatocytes during adult homeostasis and their complex plasticity upon distinct liver injuries. Cell Rep 44, 115093 (2025).

[31] Sugimoto, A. et al. Hepatic stellate cells control liver zonation, size and functions via r-spondin 3. Nature 640, 752–761 (2025).

[32] Gombin, J., Vaidyanathan, R. & Agafonkin, V. concaveman: A Very Fast 2D Concave Hull Algorithm (2020). R package version 1.1.0.

[33] Hahsler, M., Piekenbrock, M. & Doran, D. dbscan: Fast density-based clustering with R. Journal of Statistical Software 91, 1–30 (2019).

[34] Cheng, J., Smyth, G. K. & Chen, Y. Unraveling the timeline of gene expression: A pseudotemporal trajectory analysis of single-cell RNA sequencing data. F1000Res 12, 684 (2023).

[35] Cheng, J. jinming-cheng/SpNeigh: SpNeigh publication release. Zenodo (2025).

